# Perfluorooctane sulfonate affects proliferation and differentiation of pluripotent human teratocarcinoma cells

**DOI:** 10.1101/089698

**Authors:** Mina Popovic, Brett A. Neilan, Francesco Pomati

## Abstract

Perfluorinated compounds have raised concern due to their potential association with detrimental postnatal outcomes in animals and humans. We tested the effects of perfluorooctane sulfonate (PFOS) on a human pluripotent teratocarcinoma (known as NCCIT) cells as an *in vitro* prototype for developmental toxicity in mammals. NCCIT contains stem-cells able to differentiate into endoderm, mesoderm and ectoderm. We tested our cell model using a teratogenic compound, retinoic acid (RA), a cytotoxin, nocodazole (ND), and PFOS. We assayed cells proliferation, morphology and expression of stem cell and germ layer marker genes. PFOS reduced NCCIT cell proliferation in a concentration-dependent manner and induced morphological changes in cell cultures that resembled ectodermal phenotypes. A tendency towards a differentiated state in NCCIT was confirmed by real-time gene expression. PFOS triggered up-regulation of the gene nestin, indicative of ectodermal lineage differentiation, and interfered with the expression of the pluripotency stem-cell marker TERT. PFOS produced effects on both cells proliferation and differentiation, although not as severe as those observed for RA and ND, at levels that fall within the range of concentrations found in animal and human plasma. We discuss our findings in the context of possible interference of PFOS with the processes governing the early development of mammalian tissues.

## Introduction

The ubiquitous presence in the environment of perfluorinated surfactant compounds such as perfluorooctanesulfonic acid (PFOS) and perfluorooctanoic acid (PFOA) has resulted in considerable concern (Giesy and Kannan 2001; Houde et al. 2008; Jensen and Leffers 2008; Loos et al. 2007). Animal studies have indicated that PFOS and PFOA have the potential for developmental toxicity (Austin et al. 2003; Beach et al. 2006; Berger et al. 2009; Bjork et al. 2008; Giesy and Kannan 2001; Kennedy et al. 2004; Lau et al. 2004; Loveless et al. 2006). Human studies have presented epidemiological evidence linking birth defects and thyroid disease to blood serum PFOS and PFOA concentrations in the range of μg/L (Apelberg et al. 2007a; Apelberg et al. 2007b; Jensen and Leffers 2008; Melzer et al. 2010).

In this study we investigated the effects of PFOS at exposure levels relevant to animal and human sera using a stem cell-based test. Embryonic stem (ES) cells have proven to be a promising model to study developmental toxicity of chemicals (Whitlow et al. 2007). The use of ES cells, however, presents a number of challenges (see Supplementary Online Material). To avoid issues associated with the use of ES cells, we chose a human germ-cell line derived from a teratocarcinoma known as NCCIT (National Cancer Centre Immature Teratoma), which is analogue to a pluripotent ES line (Damjanov et al. 1993; Rohwedel et al. 2001; Teshima et al. 1988). Teratocarcinomas contain both undifferentiated stem-and differentiated cells that can commit to endoderm, mesoderm and ectoderm. NCCIT cells express markers of germ layers like ES cells, are capable of continuous renewal in minimal media like cancer cells (without the need of a feeder layer of primary cells) but do not show up-regulation or induction of typical tumour gene markers (Damjanov et al. 1993; Sperger et al. 2003; Taranger et al. 2005). Pluripotency in NCCIT has been proven by the expression of undifferentiated hES cells markers, including the telomerase reverse transcriptase gene (TERT) (Sperger et al. 2003; Taranger et al. 2005). Here, we aimed at using NCCIT as a stem cell model to test the effects of PFOS compared to known teratogenic and cytotoxic compounds such as retinoic acid (RA) and nocodazole (ND). We found that PFOS was able to interfere with the growth and differentiation processes of NCCIT cells.

## Materials and Methods

### Chemicals

Standard solutions were purchased from Sigma (Sigma-Aldrich, Dorset, UK). Experimental concentrations of PFOS were selected based on previously reported levels in human plasma (for details see (Apelberg et al. 2007a; Apelberg et al. 2007b; Melzer et al. 2010)). Each chemical was suspended in ethanol and stored at -20°C. RA was suspended in ethanol immediately prior to experimental testing, due to its high sensitivity to UV light and oxidizing agents.

### Cell cultures and toxicity assays

The NCCIT cell line was obtained from American Type Culture Collection (ATCC, Manassas, VA; Catalog No. CRL-2073). NCCIT cells were cultured in Dulbecco’s modified eagle medium (DMEM) (Lonza BioWhittaker, Basel, Switzerland), containing 2 mM glutamine, 4.5 g/L glucose, 10% fetal bovine serum (FBS) (Gibco-Invitrogen Corporation, Carlsbad, CA, USA), and maintained at 37°C with 5% CO_2_ in a humidified atmosphere. This work was performed using NCCIT cultures at 60% confluence.

Prior to testing, stock solutions of chemicals were diluted in DMEM (0% FBS) to obtain the maximum exposure concentration of 1 mg/L. One in three dilutions were performed in DMEM (0% FBS) to obtain 11 dilutions for each chemical ranging from 1000 to 0.017 μg/L (comprising environmentally relevant exposure levels). Test dilutions of chemicals were added to three x10^3^ cells seeded across 96-well plates (final volume 100 μL DMEM 5% FBS per well). Five untreated controls and five replicates for each concentration were prepared for each 96-well plate in each experiment and incubated for 48 h at 37°C, 5% CO_2_. Microtitre tests were performed using the CellTitre-96 AQueous One Solution Cell Proliferation Assay kit (Promega Corporation, Madison, WI). Absorbances were recorded using a microplate-reader at 490 nm (optical density-OD of the dye) and 650 nm (OD for background reference). To document cell morphology, 250 × 10 cells were seeded in 6-well plates including duplicated IC10 and IC50 concentration treatments together with untreated controls (final volume 3 mL DMEM 5% FBS), incubated for 48 h at 37°C, 5% CO_2_. Cells were then stained with trypan-blue, enumerated and microscopically photographed. More details on standard procedure for harvesting, stocking of cells, proliferation and morphological assays are provided in Supplementary Online Material.

### Effective concentration analysis

Proliferation data, expressed as percentage of MTS absorbance final values in treated cells over untreated-controls, were analysed using PriProbit (PriProbitNM, 1998-2000 Masayuki Sakuma). Concentrations resulting in 10% and 50% inhibition of proliferation (IC_10_ and IC_50_) were derived from a log-logistic symmetric fit of the concentration-effect curves, with the quality of fit to the mathematical model evaluated as an Akaike’s Information Criterion.

### RNA extraction and real-time PCR assays

Cells were seeded in 6-well plates (250 × 10^3^ cells per well) including duplicated untreated controls and duplicated IC_10_ and IC_50_ treatments with RA, ND and PFOS. Total RNA was extracted from cells 48 h after exposure to test chemicals using TRI-Reagent LS following the manufacturer’s protocol (Sigma). RNA was resuspended in RNase-free water (Invitrogen) pooling experimental duplicates for each treatment, and 1 μg of total RNA was retro-transcribed (cDNA synthesis, Marligen Biosciences, Rockville, MD, USA) using random nonamers and oligo-dT primers. Reverse transcription was performed in 3 replicates as follows: 22°C for 5 min, 42°C for 90 min and 85°C for 5 min.

Quantitative PCR targeted nestin (ectoderm), brachyury (mesoderm), alpha-fetoprotein (AFP, endoderm), and TERT (maintenance of pluripotency). Glyceraldehydes 3-phosphate dehydrogenase (GAPDH) and β-actin were chosen as housekeeping genes (see Supplementary Online Material, Table S1). Analyses were performed using QuantiTect Primer Assays and QuantiTect SYBR-Green PCR Master-Mix (Qiagen, Doncaster, Australia), optimised for efficiency by the producing company. One hundred-fold diluted cDNA samples were added to the real-time PCR reaction mix (final reaction volume of 25 μL) in three replicates for each cDNA sample. Two-step cycling was performed using the Rotor-Gene 3000A system (Corbett/Qiagen): 50°C hold for 2 min, 95°C for 2 min, 40 cycles at 95°C for 15 s and 60°C for 30 s. Expression levels and statistics for each transcript, normalised to housekeeping genes and relative to untreated cells, were obtained by comparative delta-delta-Ct method using the Relative Expression Software Tool (REST^©^) (Pfaffl et al. 2002). More details on real-time PCR protocols are reported in Supplementary Online Material.

## Results and Discussion

### Cell proliferation

RA, ND and PFOS were able to reduced NCCIT cell proliferation in a concentration-dependent manner (Fig 1). Relative to untreated cultures and after 48 h exposure, ND more strongly inhibited NCCIT proliferation compared to either PFOS or RA. IC_10_ and IC_50_ values derived for RA, ND and PFOS are summarised in Table 1. PFOS showed a lower IC_50_ than RA. Viability of cells was assayed by trypan-blue staining for RA, ND and PFOS treated cultures (data not shown), supporting the cytotoxicity assay results and suggesting induction of cell death as mechanism of reduced cells proliferation. The IC_10_ value derived for PFOS based on NCCIT cells proliferation tests falls within the range of concentrations previously reported in animal and human blood sera (Apelberg et al. 2007a; Apelberg et al. 2007b; Melzer et al. 2010). Serum collected from pregnant women and umbilical cords have been shown to contain 16.2 μg/L of PFOS (Monroy et al. 2008), while concentrations of 12.8 μg/L were obtained from general human blood serum samples (Olsen et al. 2003).

**Fig. 1.**
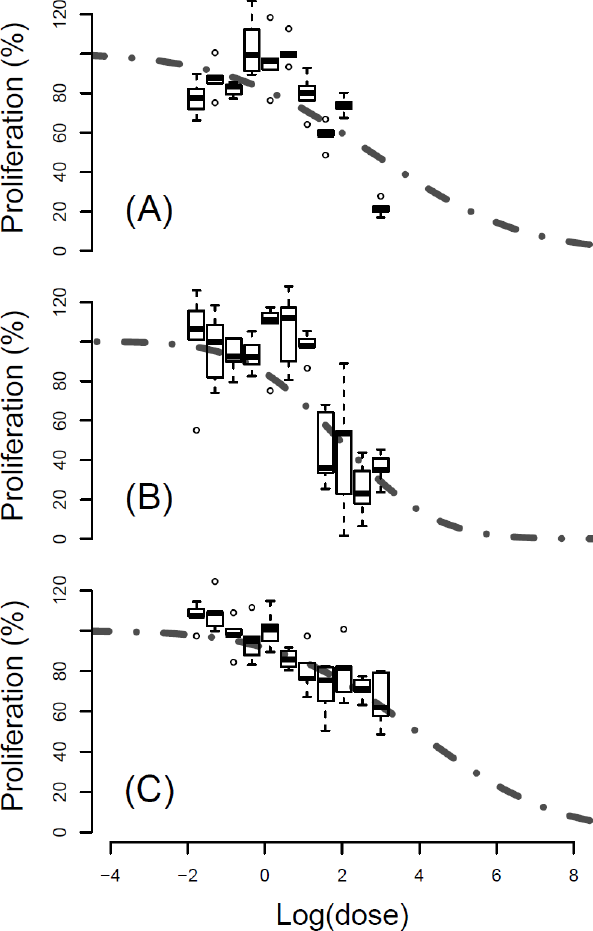
Dose-response curves for the test chemicals on the proliferation of NCCIT cells expressed as percentage of effect over control (untreated) cultures. Boxplots represent the range of experimental data-points (n=10), grey dashed lines the derived response models (see Methods). Panels depict effects of RA (A), ND (B) and PFOS (C).

**Table 1.**
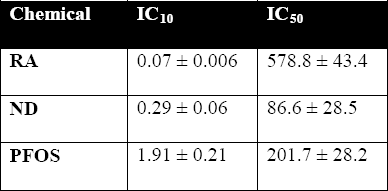
IC values (μg/L ± standard error) obtained with the NCCIT proliferation assay (48 h) for test chemicals (see Fig. 1).

| **Chemical** | **IC<sub>10</sub>** | **IC<sub>50</sub>** |
| --- | --- | --- |
| **RA** | $0.07 \pm 0.006$ | $578.8 \pm 43.4$ |
| **ND** | $0.29 \pm 0.06$ | $86.6 \pm 28.5$ |
| **PFOS** | $1.91 \pm 0.21$ | $201.7 \pm 28.2$ |

### Cell differentiation

IC_10_ and IC_50_ levels were used to assess morphological changes in NCCIT cells following treatment with RA, ND and PFOS. Cells treated with the chemicals formed smaller and flatter colonies compared to controls (Fig. 2A, Supplementary Fig. S2-S4), with multi-nucleated and more differentiated branching cells characterised by elongated cytoplasmic processes. In the IC_50_ treatments a number of detached small rounded (apoptotic) cells were evident (Supplementary Fig. S2-S4), with ND showing the strongest effects (Supplementary Fig. S3). Flattened colonies and differentiating cells with branching processes were visible in both IC_10_ and IC_50_ treatments for all chemicals (Fig. 2 B-C-D). In particular, PFOS induced changes in NCCIT cell morphology that were indicative of both differentiation and cytotoxic effects (Fig. 2D, Supplementary Fig. S4). IC_10_ exposure levels resulted in smaller and flattened colonies with differentiated cells and IC_50_ doses induced apoptosis in NCCIT, with surviving cells expressing extended cytoplasmic processes (Fig. 2D).

**Fig. 2.**
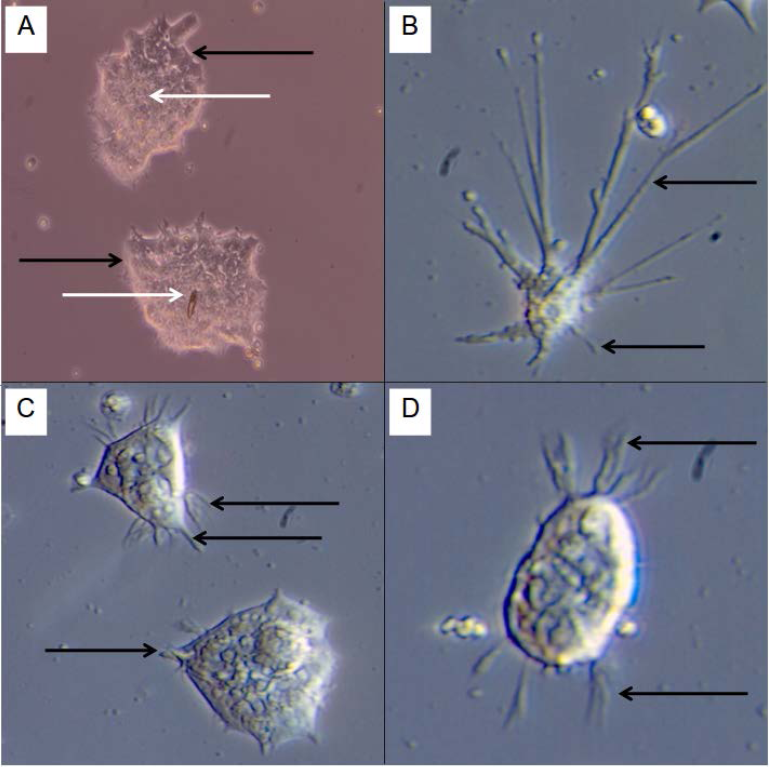
Appearance of NCCIT cells treated with the test chemicals, colour of image differs for optimal contrast. (A) Control (untreated cultures): cell colonies showing a biphasic morphology with surface cells appearing tall, predominately columnar in shape (indicated by black arrows), whereas inner core cells are mostly rounded and closely packed (indicated by white arrows). (B) IC_50_ treatment with RA: close-up of cells with long cytoplasmic processes (black arrows). (C) Close-up of IC_50_ treatment with ND: cells exhibiting elongated structures (black arrows). (D) Close-up of IC_50_ treatment with PFOS: cells with cytoplasmic processes (black arrows).

The inner mass of a NCCIT cells colony represents the equivalent of pluripotent stem cells, which normally differentiate into epithelial-like (more flattened) cells at the edges of colonies (Damjanov et al. 1993; Teshima et al. 1988). IC_10_ and IC_50_ treated cultures showed epithelial-cell features (Fig. 2-3-4 B-D) indicating a possible differentiation of NCCIT into the ectodermal germ-layer (Damjanov et al. 1993). These morphological observations were verified by the expression pluripotency/differentiation gene markers.

**Fig. 3.**
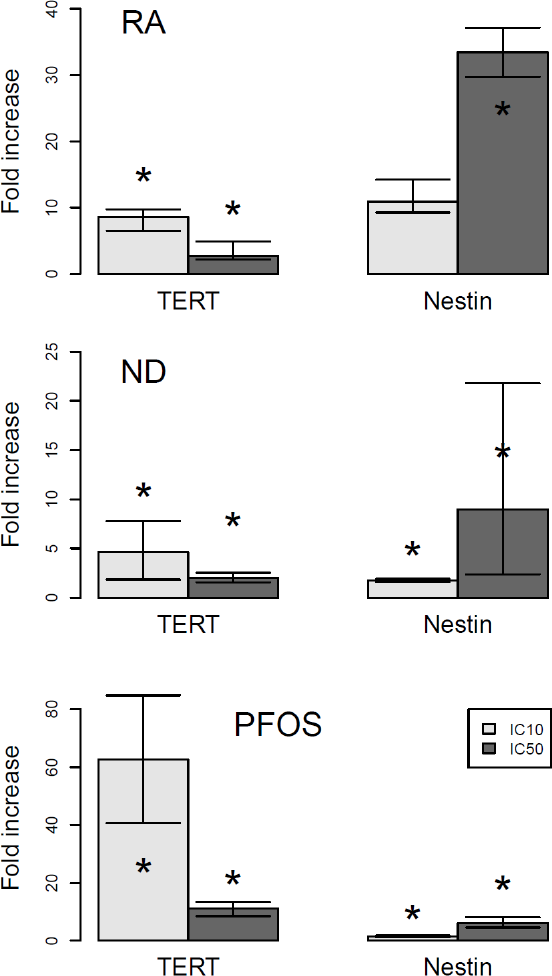
Expression profiles of TERT and nestin genes in NCCIT cells exposed to IC_10_ and IC_50_ levels of RA, ND and PFOS. Data are expressed as fold-change over the internal calibrator genes (GADPH and β-actin) and relative to control (untreated) cells. Bars on the graph correspond to the standard error of the mean (n=6). Statistical significance is reported at p < 0.05 with respect to control expression levels (*).

Exposure of NCCIT cells to RA resulted in a 8.6-fold and 2.7-fold up-regulation of TERT (pluripotency), and a 10.9-fold and 33.45-fold up-regulation of nestin (ectoderm) in the IC10 and IC50 exposures, respectively, compared to controls and relative to housekeeping genes (Fig. 3). Weaker effects on gene expression were observed for the ND treatments, in which TERT showed a 4.6-fold and 2-fold increased expression and nestin a 1.7-fold and 9fold increase in transcript accumulation for the IC10 and IC50 exposure levels, respectively (Fig. 3). PFOS was the strongest inducer of TERT mRNA expression in NCCIT, with a 62.6-fold and 11-fold up-regulation of the pluripotency marker for IC_10_ and IC_50_ exposure levels, respectively (Fig. 3). PFOS had an effect on nestin expression in NCCIT cells similar to the one exerted by ND, with 1.5-fold and 6-fold up-regulation for IC_10_ and IC_50_ exposure levels, respectively (Fig. 3).

TERT and nestin were significantly up-regulated across all levels of chemical exposure compared to the controls and relative to housekeeping genes (Fig. 3). IC_50_ doses, however, consistently resulted in reduced TERT and increased nestin genes expression compared to IC_10_ treatments (Fig. 3). Increased transcription of TERT in response to IC_10_ doses can be attributed to other functions of TERT beyond its major role in telomere maintenance (Karlseder et al. 2004) and may highlight an early response to cytotoxic effects. TERT expression in response to cytotoxic stress can enhance cell survival with an anti-apoptotic role (Cao et al. 2002; de Lange 2005). The subsequent decrease in TERT expression at increasing doses is consistent with loss in pluripotency and commitment of stem-cells to a particular lineage after exposure to teratogenic and toxic chemicals (Adler et al. 2008; Rohwedel et al. 2001). The hypothesis that NCCIT cells attempted to cope with chemical stress at low doses and then differentiate at higher exposure levels was confirmed by the detected patterns in nestin expression. An increase compared to control levels was observed at IC_10_ and IC_50_ for all test compounds (Fig. 3). Our results indicated that exposure of NCCIT cells to PFOS induced specialisation into ectodermal lineages.

### Conclusions

Our data suggest that the environmental pollutant PFOS can interfere with the functioning and differentiation processes of developing tissues *in vitro*. Given the mixed population in NCCIT, comprising both differentiated and undifferentiated cells, the tendency towards an increase in ectodermal cells after exposure to the tested chemicals may have been the result of two processes: a commitment of stem cells to ectodermal linages or a preferential selection of ectodermal cells among the population based on higher resistance towards toxins. The overall effects of PFOS were comparable to those observed for known teratogenic and cytotoxic compounds like RA and ND, at exposure levels that are relevant for risk assessment.

Giesy and Kannan (2001) have reported the global distribution of PFOS with measured concentrations in the environment and animal tissues that can exceed the IC_10_ levels derived here, although free concentration of PFOS in our *in vitro* assay (5% FBS) were probably higher than in animal sera where PFOS can bind to proteins (Jones et al. 2003). Future work should further investigate the mechanism by which PFOS can induce adverse effects on developing tissues *in vitro* and *in vivo*, considering both selection processes based on cells sensitivity in a mixed population and induction of cells differentiation. In this study, we did not detect NCCIT multi-lineage differentiation based on the expression brachyury (mesodermal marker) and AFP (endodermal marker) (data not shown). The NCCIT cell-based test presented here, however, which is accessible to any tissue-culture laboratory and can be readily standardised, can be extended by the inclusion of ectoderm expression markers and may contribute in the future to screening of pollutants for potential developmental endpoints (for more details see Supplementary online Material).

## Acknowledgements

The authors would like to thank Jason Woodhouse and Helder Marcel for technical support.

